# MicroRNA-202 *(miR-202)* controls female fecundity by regulating medaka oogenesis

**DOI:** 10.1101/287359

**Authors:** Stéphanie Gay, Jérôme Bugeon, Amine Bouchareb, Laure Henry, Jérôme Montfort, Aurélie Le Cam, Julien Bobe, Violette Thermes

## Abstract

Female gamete production relies on coordinated molecular and cellular processes that occur in the ovary throughout oogenesis. In fish, as in other vertebrates, these processes have been extensively studied both in terms of endocrine/paracrine regulation and protein expression and activity. The role of small non-coding RNAs in the regulation of animal reproduction remains however largely unknown and poorly investigated, despite a growing interest for the importance of miRNAs in a wide variety of biological processes. Here, we analyzed the role of *miR-202*, a miRNA predominantly expressed in male and female gonads in several vertebrate species. We studied its expression in the medaka ovary and generated a mutant line (using CRISPR/Cas9 genome engineering) to determine its importance for reproductive success with special interest for egg production. Our results show that miR-202-5p is the biologically active form of the miRNA and that it is expressed in granulosa cells and in the unfertilized egg. The knock out (KO) of *miR-202* resulted in a strong phenotype both in terms of number and quality of eggs produced. Mutant females exhibited either no egg production or produced a drastically reduced number of eggs that could not be fertilized, ultimately leading to no reproductive success. We quantified the size distribution of the oocytes in the ovary of KO females and performed a genome-wide transcriptomic analysis approach to identified dysregulated molecular pathways. Together, cellular and molecular analyses indicate that lack of *miR-202* impairs the early steps of oogenesis/folliculogenesis and decreases the number of large (*i.e.* vitellogenic) follicles, ultimately leading to dramatically reduced female fecundity. This study sheds new light on the regulatory mechanisms that control the early steps of follicular development and provides the first *in vivo* functional evidence that an ovarian-predominant microRNA may have a major role in female reproduction.

**Author summary:** The role of small non-coding RNAs in the regulation of animal reproduction remains poorly investigated, despite a growing interest for the importance of miRNAs in a wide variety of biological processes. Here, we analyzed the role of *miR-202*, a miRNA predominantly expressed in gonads in vertebrate. We studied its expression in the medaka ovary and knocked out the *miR-202* genes to study its importance for reproductive success. We showed that the lack of *miR-202* results in the sterility of both females and males. In particular, it lead to a drastic reduction of both the number and the quality of eggs produced by females. Mutant females exhibited either no egg production or produced a drastically reduced number of eggs that could not be fertilized, ultimately leading to no reproductive success. Quantitative histological and molecular analyses indicated that *miR-202* KO impairs oocyte development and is also associated with the dysregulation of many genes that are critical for reproduction. This study sheds new light on the regulatory mechanisms that control oogenesis and provides the first *in vivo* functional evidence that an ovarian-predominant microRNA may have a major role in female reproduction.

## INTRODUCTION

In fish, female fecundity is tightly linked to the proper completion of oogenesis in the ovary, whereby undifferentiated germinal stem cells undergo meiosis, a dramatic increase in size and ultimately form the eggs (1). Such an important differentiation processes requires coordinated interactions between the oocyte and the surrounding somatic cells (granulosa and theca cells), which together form the ovarian follicles (2). While the role of endocrine and intra-ovarian factors in these processes has been extensively studied, the regulation by the small non-coding RNAs (microRNAs) has received far less attention (3)(4).

MicroRNAs (approximately 22 nucleotides in length) play many different biological functions through the post-transcriptional regulation of protein-coding genes. In mammals, most researches that investigate the role of miRNAs in the ovary have mainly relied on *in vitro* analyses. Numerous studies that have highlighted the role played by miRNAs in ovarian development and oogenesis, as shown for miR-224 (5)(6)(7). In contrast, data documenting the role played by miRNAs in fish reproduction remain scarce and mainly rely on expression studies. In zebrafish, the expression and regulation of specific miRNAs has been associated with vitellogenesis and follicular development (8). More specifically, expression and regulation of miR-17 and miR-430b in the ovary suggest a role in follicular development and oocyte maturation. Large scale differential transcriptomic analyses were also performed during sex differentiation, gonadal development or vitellogenesis in various fish species, including rainbow trout (9), zebrafish (10)(11), Atlantic halibut (12), yellow cat fish (13), Tilapia (14)(15) and medaka (16)(17). The identification of differentially expressed miRNAs through the different stages of the reproductive cycle is consistent with a participation of miRNAs in oogenesis and spermatogenesis. Among the most abundantly expressed miRNA was *miR-202,* a miRNA known to be predominantly expressed in gonads in several vertebrate species, including frog (18), Atlantic halibut (19), human, mouse, rat (20)(21), chicken (22)(23), rainbow trout (24) and medaka (25). For protein-coding genes, such a predominant expression in gonads is usually associated with a major role in reproduction, as illustrated by many maternal-effect genes (26). There is however no functional evidence of the major role in fish reproduction of either *miR-202* or any other gonad-predominant miRNAs, although some recent *in vitro* studies reported a role for *miR-202* in mouse during spermatogenesis (27). Furthermore, it is currently unknown if such a critical feature, observed for protein-coding genes, would also be observed for miRNAs that are usually seen only as fine modulators of gene regulatory networks rather than major regulators of biological processes (28).

Here, we investigated the role of *miR-202* in fish reproduction with special attention for its role in the ovary. We thoroughly analyzed its cellular expression within the ovary and characterized the reproductive phenotype of *miR-202* knock-out (KO) fish, generated by CRISPR/Cas9 genome engineering. We showed that miR-202-5p is the predominant mature form expressed in granulosa cells. The lack of miR-202 resulted in a dramatic decrease of fertility in both male and females. In females, both egg number and quality were reduced, ultimately leading to the absence of any viable offspring. Quantitative image analysis of mutant ovaries showed that *miR-202* impairs the early steps of oogenesis/folliculogenesis. Along with this drastic phenotype, we observed the dysregulation of several key genes known for their role throughout oogenesis, as well as the dysregulation of genes, the role of which is not yet known in the ovary. Overall, this study sheds new light on the regulatory mechanisms that control the early steps of follicular development and provides the first *in vivo* functional evidence that an ovarian-predominant microRNA has a major role in female reproduction.

## RESULTS

### MiR-202-5p is highly expressed in granulosa cells

The medaka miR-202 gene harbors two mature miRNAs sequences, miR-202-5p and miR-202-3p (Fig 1A), whose expression levels were surveyed by quantitative reverse transcription TaqMan PCR (TaqManPCR) in eleven different tissues of adult fish and during embryonic development. The miR-202-5p mature form was found specifically expressed at high levels in both gonads (*i.e.* ovary and testis), in comparison to all other tissues (Fig 1B). During embryonic development, the highest expression levels of miR-202-5p were detected in non-fertilized eggs at stage 0 (st.0), thus corresponding to a maternal accumulation (Fig 1C). Lower levels of miR-202-5p were detected from the one-cell stage (st.2) onward. A slight increased expression was detected just before hatching (st.39), likely corresponding to a zygotic transcription. The miR-202-3p mature form presented similar expression profiles in adult tissues but expression levels were approximately 1500 times lower in ovary and testis, in comparison to miR-202-5p an undetectable in other tissues. During development, miR-202-3p also exhibited a profile similar to the −5p form and could be detected in unfertilized eggs at levels 500 times lower, while it could not be detected above background levels at any other developmental stages (data not shown).

**Figure 1:**
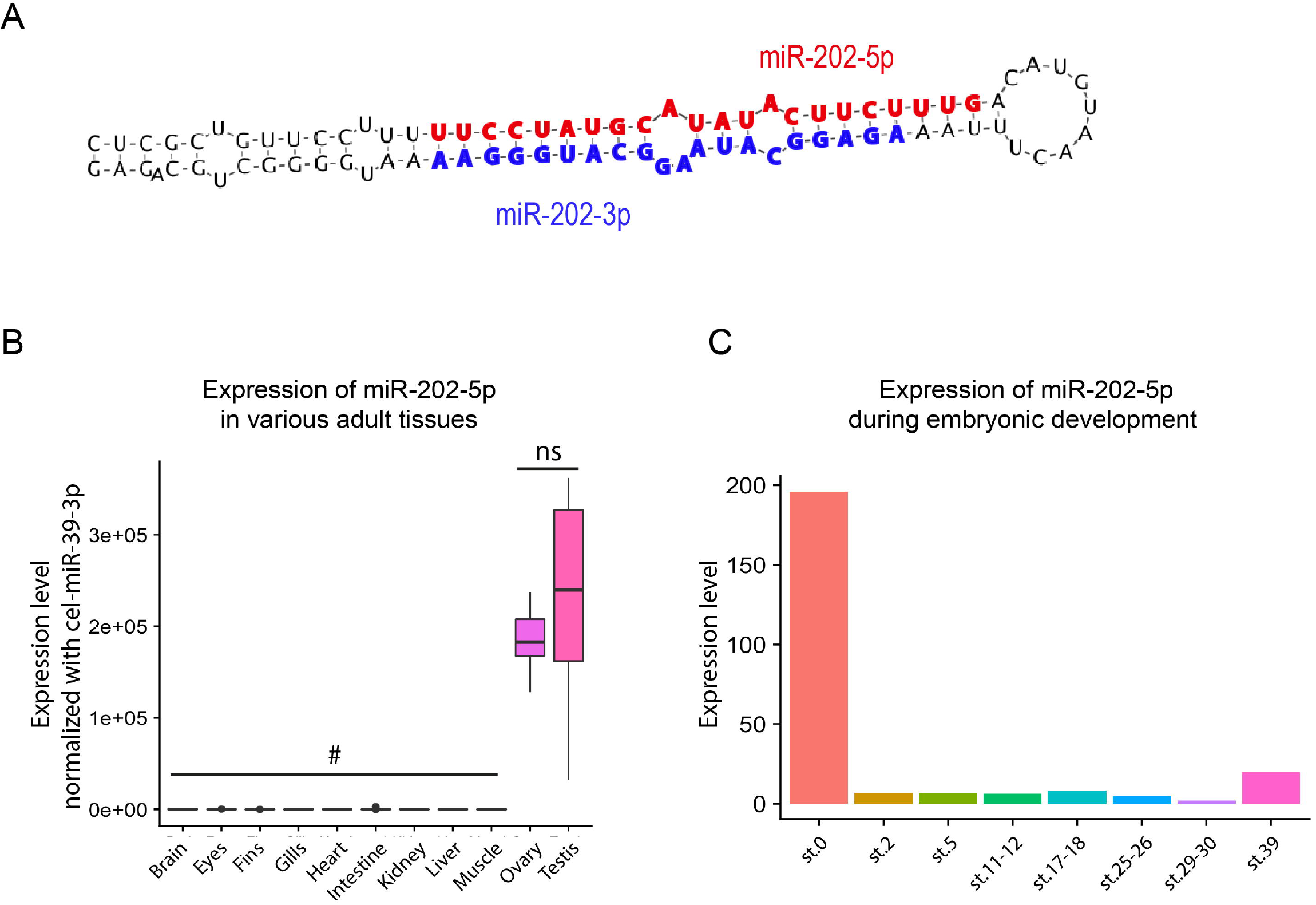
Expression profile of miR-202-5p in adult tissues and during embryonic development. **(A)** Sequence and secondary structure of the pre-miR-202. miR-202-5p is in red and miR-202-3p in blue. **(B)** Tissue distribution of miR-202-5p in eleven adult tissues obtained by TaqMan QPCR (Box plots, n=8, the ends of the boxes define the 25th and 75th percentiles; a line indicates the median and bars define the 5th and 95th percentiles). Cel-miR-39-3p was used as an external calibrator for normalization. **(C)** Expression profile of miR-202-5p during embryonic development. Pools of embryos at the same stage were used for TaqMan QPCR. *ns*, no significant difference (Student t-test). *#*expression levels not significantly different from the background signal.

The cellular expression pattern of *miR-202* was analyzed in the ovary by fluorescent *in situ* hybridization (FISH) on ovarian sections (Fig 2). Ovaries were dissected either from juvenile females (*i.e.* before the first spawning) or from adult females (*i.e.* reproductively active fishes). In both juvenile and adult ovaries, miR-202-5p was detected in follicles at all vitellogenic and post-vitellogenic stages, in granulosa cells surrounding the oocyte (Figs 2A’ and 2B’, arrowhead), but not in theca cells (Figs 2A’ and 2B’’, arrow). To visualize *sox9*-expressing cells in the germinal cradle, we performed FISH on ovaries from adult transgenic medaka *Tg*(*sox9b*::EGFP). MiR-202-5p was detected in a subset of GFP-positive cells surrounding early pre-vitellogenic follicles (primordial follicles, Fig 2B’’’). FISH performed with a specific MiR-202-3p probe (Fig 2C, D) or with a scramble control probe (S1 Fig) revealed no detectable signal above background.

**Figure 2:**
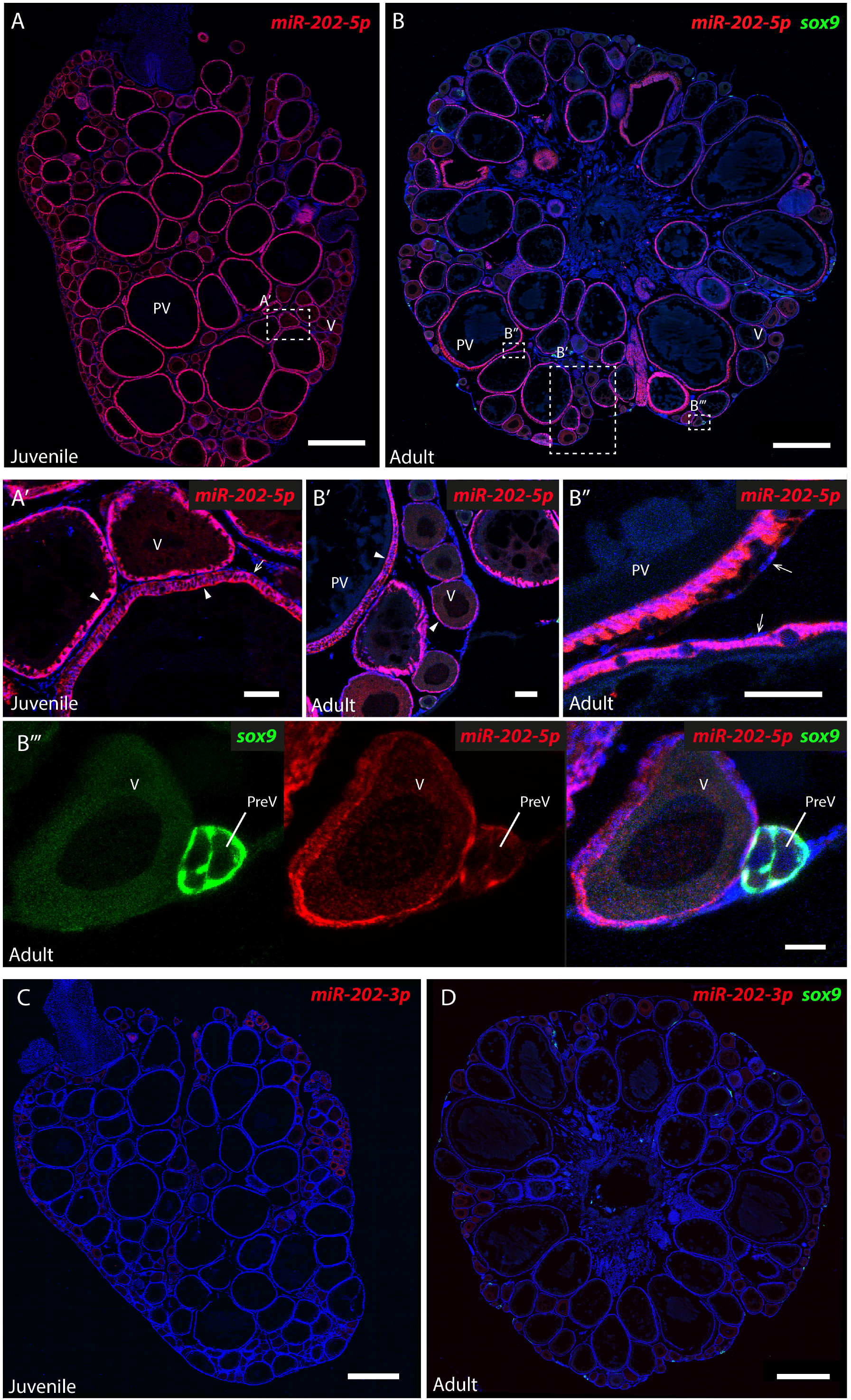
Expression of miR-202-5p and miR-202-3p in the ovary. Fluorescent *in situ* hybridizations (FISH) were performed on sections of ovaries from juvenile and adult medaka females. MiR-202-5p **(A, B)** and miR-202-3p **(C, D)** were detected with specific LNA miRNAs detection probes (in red). Ovaries of adult females were dissected from the transgenic medaka line *Tg*(*sox9b*::EGFP). The somatic GFP+ cells of the germinal cradle, including early granulosa cells, were immunodetected (in green) **(B, D)**. Nuclei are stained with DAPI (in blue). **(A’)** A magnified view of the juvenile ovarian section. **(B’, B’’ and B’’’)** Magnified views of the adult ovarian section. Mir-202-5p is detected in granulosa cells of follicles at all stages in both juvenile and adult ovary (arrowhead), but not in the theca cells (arrow). **(B’’’)** MiR-202-5p co-localize with GFP in early granulosa cells surrounding pre-vitellogenic follicles. **(C, D)** Mir-202-3p is not detected in ovaries from juvenile and adult fishes. PreV, pre-vitellogenic follicle; V, vitellogenic follicle; PV, post-vitellogenic follicles. Scale bars: 500 μm (A-D), 50 μm (A’, B’, B’’) and 20 μm (B’’’).

### *MiR-202* knock-out drastically reduces female fertility

To determine the role of miR-202 on female reproduction, we inactivated the *miR-202* gene using the CRISPR/Cas9 technology. Small insertion/deletion (INDEL) mutations were inserted in the genome, in the miR-202-3p mature sequence (Fig 3A). Fishes displaying the same INDEL (−7+3) were selected to establish a mutant fish line. In homozygotes mutants (*miR-202-/-*), the processing of the pri-miR-202 was impaired and the miR-202-5p mature form was absent (Fig 3B). The sex ratio for both heterozygous and homozygous was the same as in wild-type siblings. This mutant line was used in all further experiments.

**Figure 3:**
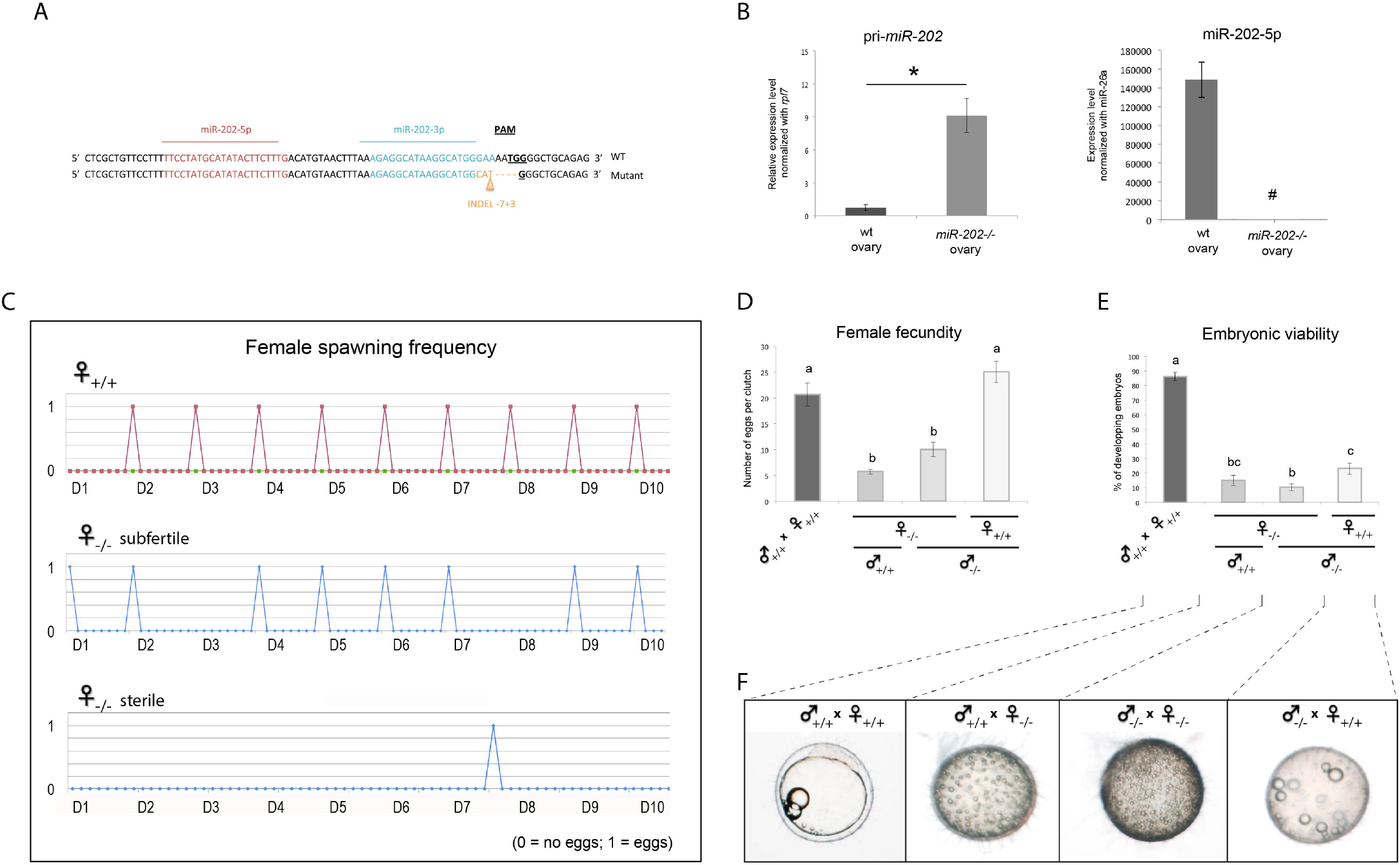
Analysis of the reproductive phenotype of *miR-202-/-* adult fish. **(A)** Genomic regions of wild-type and mutant fishes showing the INDEL mutations (−7+3) inserted in the miR-202-3p sequence using CRISPR/Cas9 genome engineering. **(B)** Expression levels of the pri-miR-202 and miR-202-5p forms in the ovaries of wild-type and homozygotes females (*miR-202-/-*) were measured by QPCR. **(C)** Spawning frequency of mutant and wild-type females were monitored during ten days. Subfertile females (irregular spawning) and sterile females (only one spawning) represent respectively 85% and 15% of all analyzed females. **(D)** Number of eggs per clutch spawned by wild-type or subfertile females when mated with wild-type or mutant males. **(E)** Embryonic viability of spawn eggs measured by the percentage of eggs that are fertilized and that develop correctly. **(F)** Representative pictures of eggs from the different crosses of wild-type and mutant males and females. Mean values (± SEM) are displayed on the graphs. * indicates expression levels that are significantly different (Mann Whitney test, *p* < 0.05). Different letters indicate a significant difference, as determined by a one-way ANOVA test (Tukey’s post hoc test). *#*expression levels not significantly different from the background signal.

We thoroughly analyzed the reproductive phenotype of *miR-202-/-* adult females. The frequency of spawning was analyzed during ten consecutive cycles (Fig 3C). Two categories of mutant females were distinguishable by different reproductive phenotypes. The mildly affected females (subfertile females), which represented 85% of all mutant females analyzed, displayed an irregular frequency of spawning compared to wild-type females that spawned every day within one hour of the onset of the light. The most strongly affected females (sterile females), which represented 15% of all mutant females analyzed, spawned only once in ten days. Further analysis of the quantity of spawned eggs (female fecundity) revealed that subfertile females spawned a significantly reduced number of eggs (5.8 eggs per clutch on average) compared to wild-type females (18.8 eggs per clutch on average, Fig 3D). In addition, the viability of embryos (*i.e.* capability of eggs to be fertilized and to develop correctly) was dramatically reduced when subfertile females were outcrossed with wild-type males (Fig 3E). Similar results were obtained when subfertile females were mated with mutant males. Conversely, when mutant males were outcrossed with wild-type females, the viability of embryos was also drastically affected compared to wild-type siblings. Further analysis of siblings revealed that, eggs originating from subfertile females fertilized by either a wild-type or a mutant male could not be fertilized, while eggs originating from wild-type females fertilized by a mutant male were fertilized but arrested during the first cleavage stages of embryonic development (st.4-5, Fig 3F). The overall reproductive success of mutant females was therefore reduced to 0.83 viable eggs per clutch for subfertile females (*i.e.* mutant females that exhibited irregular spawning, reduced fecundity and low embryonic survival) and to zero viable egg for sterile females that exhibited the most severe phenotype, while wild-type female exhibited an average of 15.3 viable eggs per clutch. To maintain the mutant fish line (INDEL −7+3), heterozygous fishes were thus systematically outcrossed with wild-type fishes at each generation (see Material and Methods). Another mutant line harboring another INDEL mutation (−8) was used to confirm this reproductive phenotype but this line was not used for further histological and molecular phenotyping analyses (data not shown). Altogether, our results indicate that *miR-202* is required for the formation of functional gametes in both female and male gonads.

### *MiR-202* knock-out affects the early steps of follicular growth in juvenile females

We further analyzed the role of *miR-202* during oogenesis in juvenile females (*i.e.* before the first spawning). To this aim, the follicular content of wild-type and mutant ovaries were analyzed and compared. For each ovary, the number and size of follicles were determined on the median ovarian section. Nuclei of somatic cells, including ganulosa cells surrounding each oocyte, were stained with DAPI, which allowed delineating all follicles and automatically individualizing them, using a computational automatic segmentation procedure (Fig 4A). The section area, the mean follicle diameter and the number of follicles were determined (Fig 4B). In *miR-202-/-* ovarian sections, the mean follicular diameter was significantly reduced (70 μm), as compared to wild-type ovarian sections (80 μm), and the number of follicles appeared higher (500 μm) as compared to wild-type (300 μm), although not significantly. These data suggest an impairment of the early follicular growth in juvenile mutant ovaries.

**Figure 4:**
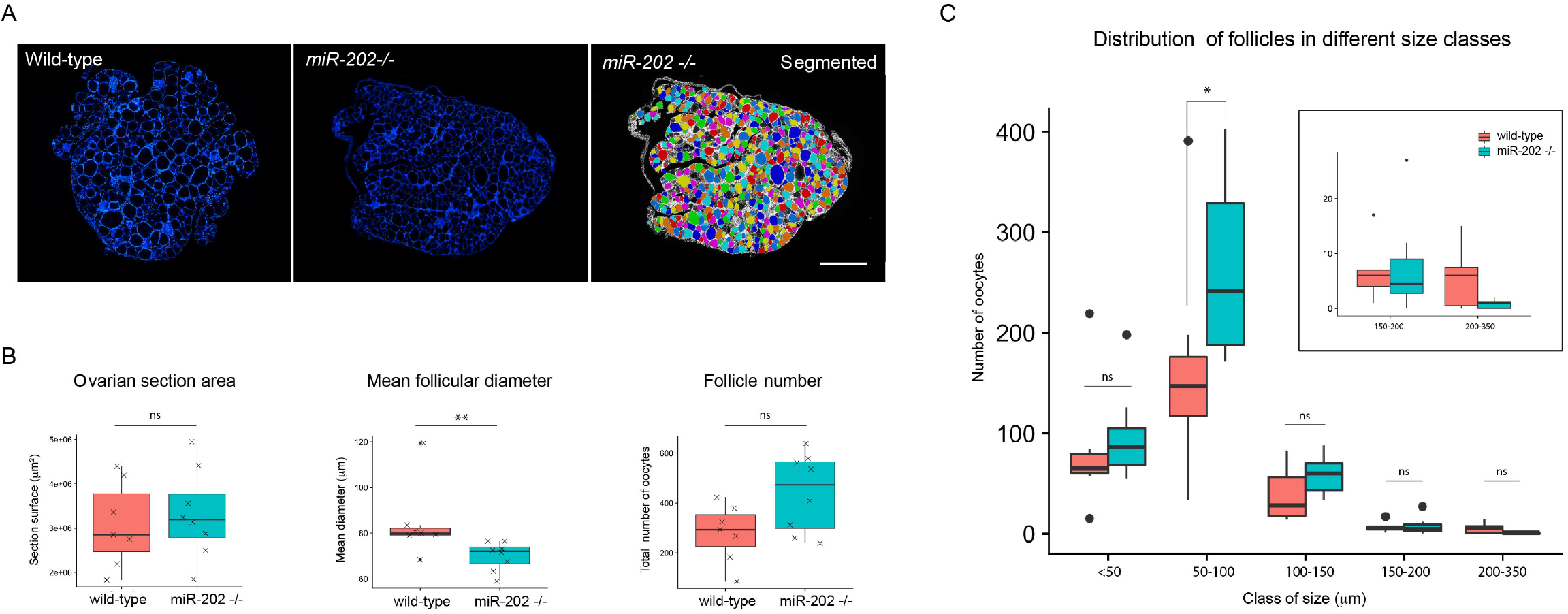
Quantitative image analysis of *miR-202-/-* juvenile ovaries. **(A)** Median sections of ovaries from juvenile wild-type and *miR-202-/-* fishes. All nuclei are stained with DAPI (in blue). Image sections were automatically segmented. **(B)** Section area, mean follicle diameter and follicle number automatically determined. **(C)** Size distribution of follicles on sections. The number of small follicles (50-100 μm) is significantly higher compared to wild-type, while the number of follicles in the larger diameter class (200-350 μm) tends to decrease. Box plots are displayed on graphs for wild-type (n=7) and *miR-202-/-* (n=8) juvenile females (in red and green, respectively). The ends of the boxes define the 25th and 75th percentiles; a line indicates the median and bars define the 5th and 95th percentiles. Individual values are shown for the graphs of panel B. Asterisks indicate significant differences (* *p* < 0.05 and ** *p* < 0.01, Mann Whitney test). Scale bar: 500 μm.

The size distribution of follicles on sections was thoroughly analyzed in mutant and wild-type ovaries. Follicles were classified according to their diameter into five different classes (<50 μm, 50-100 μm, 100-150 μm, 150-200 μm and 200-350 μm) (Fig 4C). Comparison of the resulting profiles revealed different size distributions profiles in mutant and wild-type ovaries. In mutant ovarian sections, the number of small follicles appeared higher compared to wild-type, with a significant increased in the 50-100 μm size class, whereas the number of follicles in the larger diameter class (200-350 μm) tended to decrease, although not significantly. This clearly suggests an important defect of the early follicular growth in juvenile *miR-202-/-* females, leading to an accumulation of small and medium oocytes (*i.e.* pre-vitellogenic and early-vitellogenic oocytes) to the detriment of larger late-vitellogenic and maturing follicles.

### *MiR-202* knock-out decreases the number of growing follicles in subfertile adult females

The role of *miR-202* during oogenesis was analyzed at the adult stage (reproductively active fish), in both subfertile and sterile females (Fig 5A). Section areas, mean follicular diameters and numbers of follicles were measured on median sections. The follicular size distribution was also analyzed. Follicles were classified according to their diameter into three different size classes based on their diameter: small (<100 μm), medium (100-400 μm) and large (400-1200 μm). In sterile females, ovaries displayed a significant decrease of the mean follicular diameter (170 μm) in comparison to wild-type ovaries (220 μm), but similar numbers of follicles were found in both cases. Consistently, ovarian sections were significantly reduced in mutant compared to wild-type (Fig 5B). The analysis of the size distributions also revealed distinct profiles in wild-types and mutants. The number of large follicles (>400 μm) was significantly reduced in mutant, while small follicles (<100 μm) tended to accumulate, although not significantly (Fig 5C). These observations suggest a strong impairment of the early steps of the follicular growth in sterile females, similarly to juvenile mutant females analyzed during the first reproductive cycle (see above).

**Figure 5:**
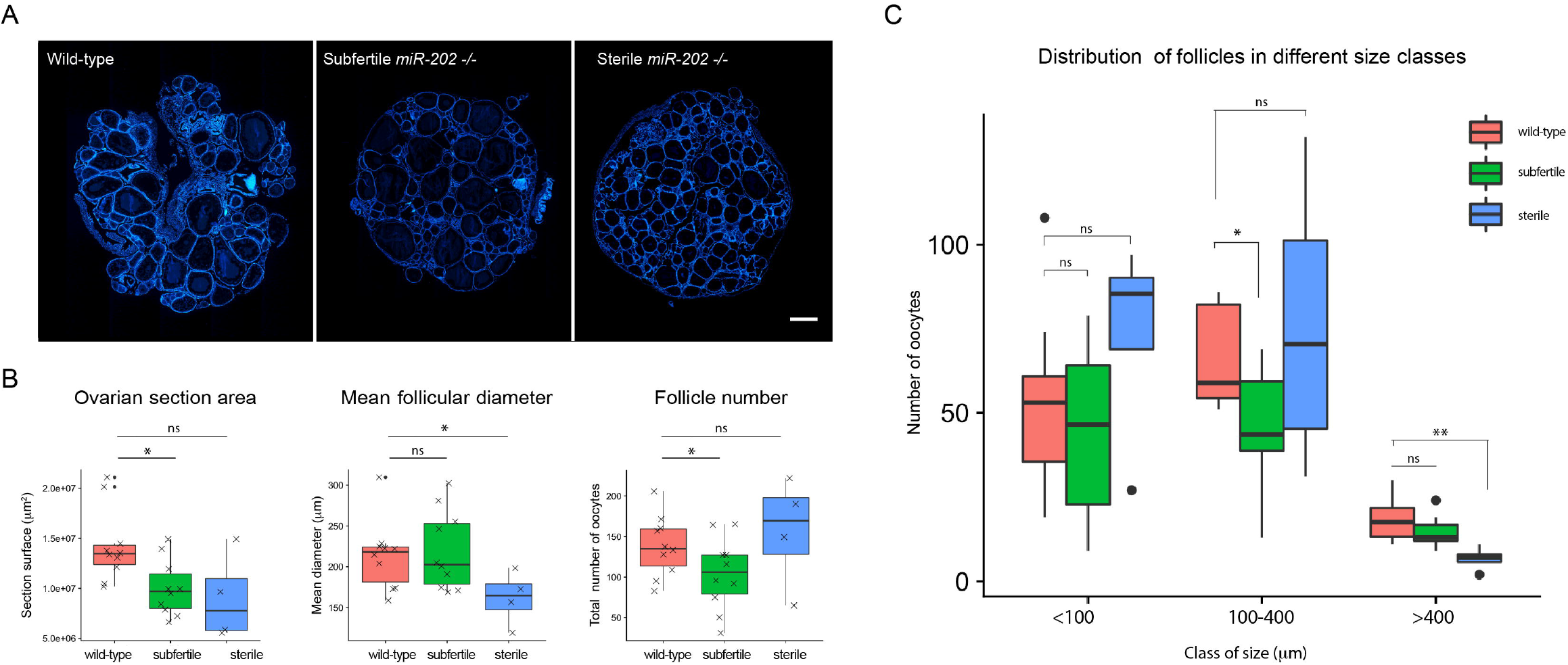
Quantitative image analysis of *miR-202-/-* adult ovaries. **(A)** Median sections of ovaries from wild-type, sterile and subfertile mutant females. All nuclei are stained with DAPI (in blue). Image sections were automatically segmented. **(B)** Section area, mean follicle diameter and follicle number automatically determined. **(C)** Size distribution of follicles on sections. In ovaries from sterile females, the number of large follicles (>400 μm) is significantly reduced, while small follicles (<100 μm) tend to accumulate. In the ovary from subfertile females, the number of medium follicles (100-400 μm) is significantly reduced, but small size follicles (<100 μm) do not accumulate. Box plots are displayed on graphs for wild-type (n=10), *miR-202-/-* subfertile (n=10) and *miR-202-/-* sterile (n=4) adult females (in red, green and blue, respectively). The ends of the boxes define the 25th and 75th percentiles; a line indicates the median and bars define the 5th and 95th percentiles. Individual values are shown for graphs of panel B. Asterisks indicate significant differences (* *p* < 0.05 and ** *p* < 0.01, Mann Whitney test). Scale bar: 500 μm.

In subfertile females, which exhibited a milder reproductive phenotype (*i.e.* occurrence of spawning but a reduced number of eggs and reduced egg survival), ovarian sections displayed a significantly reduced number of follicles. This was associated with a significant reduction of ovarian section areas (Fig 5B). No modification of the mean follicular diameter was however observed, suggesting that although fewer follicles were engaged into growth, they were all able to reach their correct final size. This was supported by the follicular size distribution profiles, which showed a significant and important reduction of the number of medium-size follicles (100-400 μm in diameter) in mutant compared to wilt-type, and no accumulation of small-size follicles (<100 μm in diameter, Fig 5C). These data likely reflect a milder effect of the *miR-202* deficiency on follicular growth in subfertile adult females compared to sterile fishes, indicating a partial recovery of the phenotype at this stage.

### Genome-wide analysis reveals major dysregulation of key ovarian genes

To get further insight into the molecular pathways that are affected in absence of miR-202, we performed a genome-wide transcriptomic analysis on ovaries of wild-type and *miR-202-/-* females. This analysis was conducted on ovaries from juvenile females, before the first spawning, when ovaries display a more homogenous follicle population (from 30 to 350 μm in diameter) compared to ovaries from reproductively active adult fish (from 30 to 1100 μm in diameter, see Figs 4 and 5). We identified 52 differentially expressed genes, including 11 genes that were up-regulated in mutant and 41 genes down-regulated in mutant (S1 Table). The most differentially expressed genes were validated by QPCR (*wnt2bb*, *wnt4a*, *klhl23, setd4, npr1b and* srgap3) along with other major genes involved in fish oogenesis selected from the literature (*cyp19a1a*, *cyp17, gsdf, inh, foxl2b, sycp3, foxl3, Sox9b, olvas*). Results are shown on figure 6 and are described bellow.

**Figure 6:**
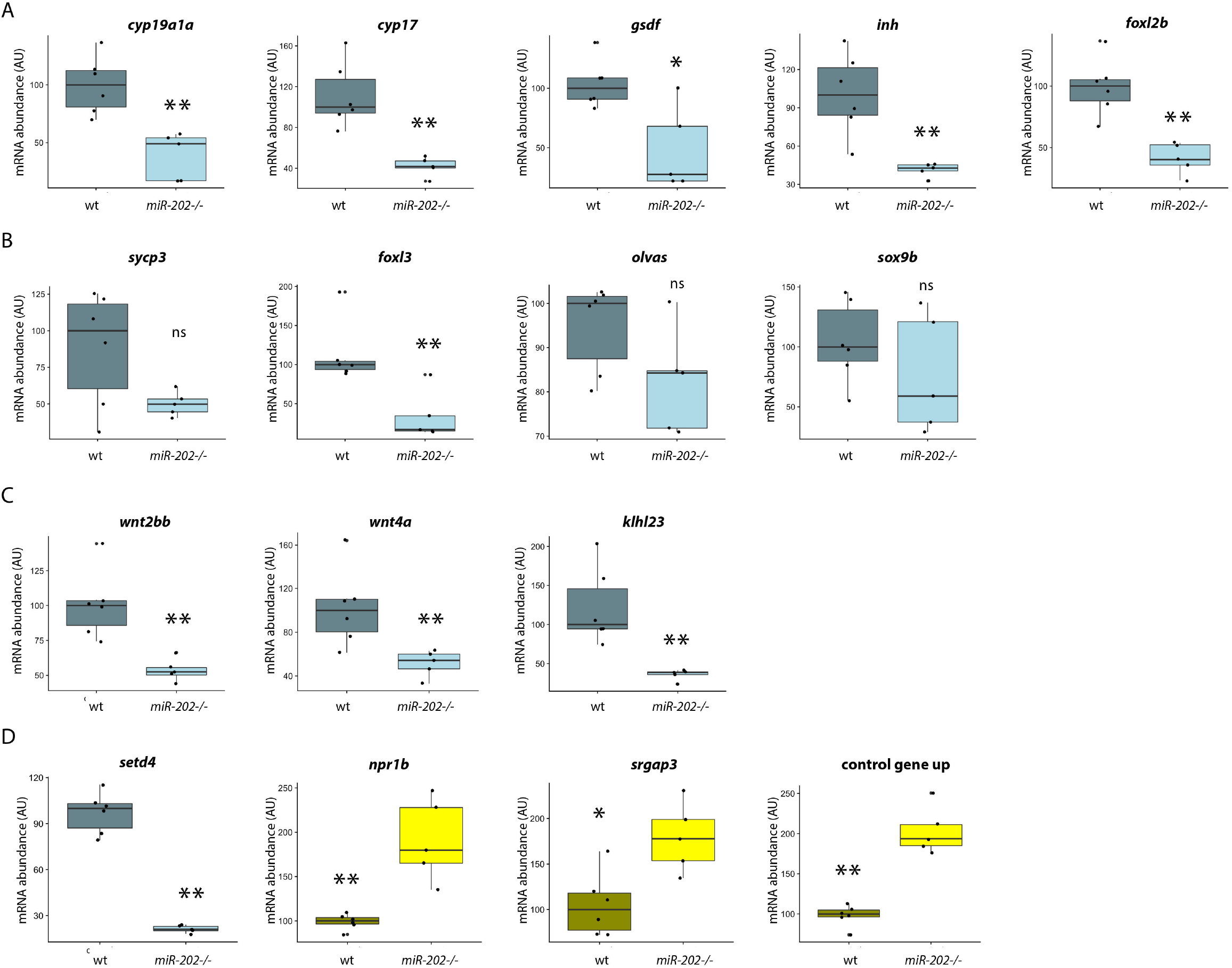
mRNA expression profiles of ovarian genes in *miR-202 -/-* females. Gene expression levels in wild-type and ovaries from juvenile females were obtained by QPCR. (A) Genes expressed in the somatic germ-cell supporting cells. **(B)** Genes expressed in primary oocytes within the germinal cradle. **(C)** Genes that belong to the WNT and KELCH families. **(D)** Dysregulated genes that have never been previously described as molecular players of oogenesis. Expression levels were measured in triplicates. Box plots are displayed on graphs for wild-type (n=6) and *miR-202-/-* (n=5) juvenile females. The ends of the boxes define the 25th and 75th percentiles; a line indicates the median and bars define the 5th and 95th percentiles. Individual values are shown. The *rpl7* gene was used for normalization. Asterisks indicate significant differences (* *p* < 0.05 and ** *p* < 0.01, Mann Whitney test).

### Down-regulation of known genes expressed in the somatic germ-cell supporting cells (Fig 6A)

Expression analysis revealed a marked expression decrease of two genes encoding key steroidogenic enzymes (*cyp19a1a* and *cyp17*). *Cyp19a1a* (also known as aromatase) is an ovarian-specific steroidogenic enzyme that mediates estradiol (E2) production and therefore plays a major role in both sex differentiation and vitellogenesis (Lubzens et al. 2010, for review). An even more pronounced under-expression was observed for *cyp17* that acts upstream of *cyp19a1a*. Consistently, data obtained from microarray analysis also revealed the down-regulation of the *steroidogenic acute regulatory* gene (*star*, S1 Table). *Star* is involved in the cholesterol shuttling across the inner mitochondrial membrane, which is a rate-limiting step of the steroidogenesis (29)(30). A decrease in *star* expression indicates an overall decrease in ovarian steroidogenic production.

We also observed the down-regulation of *gsdf* and *inhibin*, two members of the TGFβ family, in the ovary of mutant females. *Gsdf* is known to be expressed in granulosa cells and to be involved in the maintenance and maturation of these cells (31)(32). *Inhibin* consists of a unique α subunit and an activin β subunit. In the zebrafish ovary, members of the activin-inhibin-follistatin system exhibit dynamic changes in expression during folliculogenesis and the α subunit (*inha*) is expressed during vitellogenesis, especially during final oocyte growth (*i.e.* prior to final oocyte maturation)(33).

Finally, QPCR analysis revealed a significant down-regulation of the transcription factor *foxl2b*, the only *foxl2* copy retained in medaka after teleost-specific genome duplication (34). In medaka, *foxl2b* is a female-specific gene expressed in somatic cells surrounding germ-cells in the germinal cradle. At later stages, *foxl2b* continue to be expressed in granulosa cells and in a minority of theca cells in pre-vitellogenic and vitellogenic follicles, but expression ceased in post-vitellogenic follicles (35)(36). In mouse, *foxl2* is known as a key regulator of oogenesis and plays a critical role in ovarian differentiation (37)(38)(39).

### Down-regulation of known genes expressed in the primary oocyte within the germinal cradle (Fig 6B)

We analyzed the expression of *sycp3, foxl3* (a *foxl2*-relative)*, sox9b* and the medaka *vasa*-like gene (*olvas*)*. Sycp3 and foxl3* are both expressed in germ cells in the ovary and are both involved in the first meiotic division (4)(40). *Olvas* is known as a specific marker of early germ cells, which are nested in the cords of *sox9b*-expressing cells, referred to as the germinal cradle (41)(4). QPCR analysis revealed a down-regulation of *sycp3* and *foxl3* in the ovary of mutant females, although the diminution of *sycp3* expression is not significant. In contrast, the expression of *olvas* and *sox9b* was similar in wild-type and mutant ovaries. Together these observations suggest that germinal cradles of mutant ovaries are subjected to a reduced meiotic activity of primordial oogonia, while the mitotic activity of germline stem cells is likely to be maintained.

### Down-regulation of other genes that belong to the WNT and KELCH families (Fig 6C)

Among the most differentially expressed genes were several other genes coding for proteins of the WNT signaling pathway, including *wnt2bb* and *wnt4a*. In ovary of mutant females, QPCR revealed a clear and significant down-regulation of *wnt2bb* and *wnt4a*. *Wnt4a* is expressed in fish ovary, while the expression of *wnt2bb* remains uncharacterized to date (42). The microarray analysis also revealed a dramatic down-regulation of *klhl23*, a member of the KELCH Like Family. KELCH proteins were initially characterized due to their important role in oocyte-somatic cells communication during drosophila oogenesis. Several members of this large family (over 30 members) are known to be expressed in the ovary of several vertebrates even though they remain poorly studied (43).

### Identification of novel molecular players of oogenesis (Fig 6D)

Finally, transcriptomic analysis revealed three other differentially expressed genes (*setd4*, *npr1b* and *srgap3*) that have never been previously described as molecular players of oogenesis. The *set domain-containing protein 4* (*setd4*) gene displayed a significantly reduced expression in ovary of mutant females. *Setd4* encodes for a recently characterized methyltransferase involved in breast cancer cell proliferation (44). While the role of *setd4* in the ovary remains unknown, a recent report demonstrated its role in the regulation of cell quiescence during the diapause of artemia embryos (45). The *natriuretic peptide receptor 1b* gene (*npr1b*) exhibited a marked over-expression. Although the role for *npr1b* in oogenesis has never been reported, the mouse *nrp2* gene has already been detected in granulosa cells and shown to maintain meiotic arrest of oocytes in mouse ovary (46). The *slit-robo Rho GTPase activating protein 3* (*srgap3*) gene displayed a milder, yet significant, over-expression. Along with a well-established function in axon guidance, several studies suggested that the SLIT/ROBO pathway could also have important functions in the reproductive system (47). These results suggest that *setd4*, *npr1b* and *srgap3* are indeed novel molecular players of oogenesis. Given the greater number of down-regulated genes, in comparison to up-regulated genes, a non-annotated gene was selected for QPCR as a control and the profile observed on microarrays could be validated.

## DISCUSSION

Most of our current understanding of the regulatory mechanisms of oogenesis in fish is due to several decades of research on hormones, secreted factors or intrinsic signaling pathways. However, miRNAs are well known for their role in many physiological processes through post-transcriptional gene regulations. Here, we showed that this fundamental class of molecules, endowed with pleiotropic functions, is also necessary for oogenesis in medaka ovary. In particular, we showed that *miR-202* plays a key role in follicular recruitment and growth, and ultimately in the female reproductive success, as shown by the severe phenotype of *miR-202* KO fishes.

### Mir-202-5p is the biologically active mature form in the ovary

Our present results show that miR-202-5p is the predominant mature form processed from *miR-202* gene, while the miR-202-3p form is detected at much lower levels. This is consistent with existing data in which one miRNA form is biologically active, while the other is degraded. More specifically, our ISH experiments reveal that miR-202-5p is highly expressed in the somatic granulosa cells surrounding the oocyte. Consistently, recent transcriptomic studies in zebrafish reported either an increased expression of miR-202-5p during folliculogenesis or an increased expression in developing oocytes and unfertilized eggs (48)(49). Yet, it has also been claimed that *miR-202* could by expressed only in early medaka oocytes and absent at later stages, based on ISH experiments (16). Since our ISH did no reveal any expression of miR-202-5p above background levels in early oocytes, we presume that it is more likely accumulated during vitellogenesis in vitellogenic and/or post-vitellogenic oocytes, as suggested by the detectable expression in unfertilized eggs,

### *Mir-202* is required for both male and female reproductive success, but not for sex determination

Here, we observed that the inactivation of *miR-202* in medaka does not lead to any modification of the sex ratio, since mutant siblings give rise to 50% of adult males and 50% of adult females. This rules out the possibility of a key role of *miR-202* in sex-determination. It should however be stressed that the *miR-202* KO results in the sterility of both females and males. More particularly, when *miR-202* KO males were crossed with wild-type females, embryonic viability was dramatically reduced since most embryos were arrested during the first cleavage stages (st.4-5). To our knowledge, this phenotypic defect of male reproduction as never been reported before in fish, and it clearly indicates that *miR-202* is also required for spermatogenesis. This is in agreement with results obtained after *miR-202* KO in cultured spermatogonial stem cell in mouse, indicating a role in the control of the cellular proliferation/differentiation balance (27). Further studies are now required to unravel the precise role of *miR-202* in fish testis during spermatogenesis.

### Females lacking *miR-202* produce eggs that cannot be fertilized

In this study, we paid special attention to the severe phenotype displayed by *miR-202* KO females. The overall reproductive success of mutant females was reduced to 0.83 or, even more, to O viable eggs per clutch, compared to 15.3 viable eggs per clutch for wild-type females. This phenotype is the consequence of both a reduced fecundity (*i.e.* reduced number of spawn eggs) and reduced egg viability. Mutant females indeed produce a large proportion of eggs that cannot be fertilized, subsequently resulting in no embryonic development. During folliculogenesis, the somatic cells that surround the growing oocyte (granulosa and theca cells) contribute to the formation of the chorion, and the micropyle, which is primordial for the egg fertilization (50). It is thus likely that *miR-202*, which is expressed in granulosa cells throughout folliculogenesis, contributes to the formation of a functional chorion and micropyle. However, we cannot totally exclude an additional role of *miR-202* in oocytes during the growth and maturation steps, given the possible accumulation of *miR-202* inside the oocyte.

### *MiR-202* deficiency severely affects early folliculogenesis and female fecundity

The very low fecundity of *miR-202* KO females strongly indicates that *miR-202* plays an important role in the oogenesis processes in the ovary, which was further investigated. One of the most remarkable features observed in mutant ovaries was the abnormal increased number of pre-vitellogenic follicles. Along with this phenotype, we observed a reduced expression of *gsdf* (a member of the TGFb family) and of the *foxl2* transcription factor. Both *gsdf* and *foxl2* are expressed in granulosa and are well known for their role as key regulators of early oogenesis/folliculogenesis (34)(51)(52). Downregulation of these factors thus confirms that *miR-202* likely controls the early development of granulosa cells (*i.e.* granulosa proliferation and/or differentiation) through the regulation of *gsdf* and *foxl2*, and ultimately leads to an impaired folliculogenesis. Such hypothesis is also supported by the down-regulation of downstream key genes, including steroidogenic genes (*star*, *cyp19a1a*, and *cyp17*) and the TGF-b family member *inhibin*. The formers are involved in estrogen synthesis in ovarian follicles, while *inhibin* plays a critical role during follicular development in zebrafish (53). Further analysis of the proliferation and differentiation states of granulosa cells are needed to fully understanding of the role of *miR-202* during folliculogenesis.

Of particular interest is also the down-regulation of *gsdf* in *miR-202* mutant ovaries. Along with its well-established key role in the male-determination in medaka, the GSDF factor is also thought to be necessary for normal ovarian development (54)(31). Indeed, *gsdf* inactivation not only impairs male differentiation, but also leads to a severely reduced female fecundity up to a total infertility for the most severe cases. In such mutants, the number of pre-vitellogenic follicles is abnormally high (31). Except the male-determination phenotypic defect, many features of the *gsdf* mutant are very similar to that observed for *miR-202* KO females, including the pre-vitellogenic follicles accumulation. We hypothesize that *miR-202* promotes *gsdf* expression in the ovary and boost its action on follicular development. In any case, such hypothesis remains to be confirmed and molecular mechanisms underlying the interaction between miR-202-5p and *gsdf* remain to be identified.

### *Mir-202* knock-out impairs female meiosis

The germinal cradle is composed somatic *sox9b*-expressing cells that surround mitotic and meiotic germ cells, as well as primordial oogonia barely engage into folliculogenesis (4). Our results show that *miR-202* is expressed in the germinal cradle, in a subset of *sox9b*-expressing cells surrounding primordial oogonia, which together likely correspond to primordial follicles. Interestingly, no modification of *sox9b* expression was observed in mutant ovaries, and previous studies in mouse have shown that *sox9* regulates the expression of *miR-202* in granulosa cells (55). It is thus possible that *miR-202* acts downstream of *sox9b* in medaka as well. However, molecular mechanisms that regulate the expression of *miR-202* in follicles at later stages, in vitellogenic and post-vitellogenic follicles, remain to be determined since *sox9b* is not expressed at these stages.

We also observed, in juvenile ovaries, a down-regulation of the early meiotic oocyte markers *foxl3* and, to a lesser extent *sycp3*, even though no *miR-202* expression was detected in oocytes at this stage. Cellular interactions between germinal and somatic cells are thought to be important and to be mediated by several secreted factors (56)(57)(43). Consequently, any early defect of oocytes is likely to occur in the context of granulosa dysfunction (discussed above), including the onset of meiosis. One possibility is that *miR-202* depletion in the granulosa affects oocyte-somatic communications, as suggested by the down-regulation of KELCH proteins that have important functions in oocyte-somatic cell interactions in drosophila (43). This hypothesis is also supported by the dysregulation of the *npr1b* receptor and the intracellular effector *srgap3* of the SLIT/ROBO signaling pathway, which could both also mediate cellular communications in the ovary (46)(47).

### *Mir-202*, a major player rather than a fine modulator

As discussed above, a drastic reduction of the reproductive success was observed for both male and female medaka lacking *miR-202,* including a reduced female fecundity and the production of poor quality eggs that cannot be fertilized. This phenotype was confirmed using two mutant lines bearing different INDELs obtained with the same guide RNA. It is generally considered that the purview of miRNAs is more likely the maintenance of regulatory networks, by fine-tuning gene expression, rather than the establishment of key regulatory networks for developmental decisions or core physiological processes. This concept is supported by the fact miRNAs KO are commonly associated with “modest” phenotypic effects, which are strongly exacerbated only under particular condition, as for example manipulations, stresses or disease conditions (28). This is in contrast with the drastic reproductive phenotype observed here for *miR-202* KO fishes. It is however possible that *miR-202* modulates a large networks of targets, which would have a synergistic effect on key regulatory pathways for folliculogenesis, such as *gsdf* (as discussed above), and ultimately leads to subfertile or sterile females. Nevertheless, the phenotypes observed in *miR-202-/-* fish are among the most severe observed after miRNA KO, and it would be very informative in the future to determine whether the drastic *miR-202* KO phenotype is an exception in the ovary.

### A new look at oogenesis

Surprisingly, the transcriptomic analysis performed in juvenile females, during the first reproductive cycle and before the occurrence of the first spawning, did not result in the identification of a large number of differentially expressed. It should however be stressed that among de differentially expressed genes were many genes that are crucial for steroidegenesis and reproduction, such as *star* and members of the *wnt* family (as discussed above). Despite these usual suspects, the transcriptome analysis also shed light on other genes such a genes of the *kelch* family that are less studied but believed to play an important role in oogenesis based on existing data in other animal species (58). Finally, the identification of genes that were previously not known to participate in oogenesis, such as *setd4*, *npr1b* and *srgap3*, could shed a new light on our understanding of this complex and coordinated biological process. This begs for further investigations that will greatly benefit from *miR-202-/-* fishes as a novel biological model.

## Conclusion

In summary, our results show that *mir-202* is a key miRNA involved in the regulation of follicular recruitment and growth. This provides the first functional evidence that microRNAs are necessary for the female reproductive success and in particular the regulation of female fecundity. Furthermore, the present study shed new light on the regulatory mechanisms that control the early steps of follicular development, which remain poorly understood to date. A further systematic *in vivo* functional analysis of other ovarian-predominant miRNAs should greatly increase our knowledge on the overall role of miRNAs in oogenesis and female fecundity in fish.

## Materials and methods

### Ethics statement

All experimental procedures were conducted in strict accordance with the French and European regulations on animal welfare recommendations and were approved by the INRA LPGP Animal Care and Use Committee.

### Medaka breeding and tissues collection

Adult medaka (*Oryzias latipes*) from the CAB strain and adults of the *Tg(sox9b::EGFP)* medaka line were raised at 26°C. Fishes were raised under a growing photoperiod regime until 3 months post-fertilization (12h light/ 12h dark) and under a reproduction photoperiod regime after 3 months post-fertilization (14h light/10h dark). For QPCR analysis, eleven different tissues/organs (brain, eyes, fins, gills, heart, intestine, kidney, liver, muscle, ovary and testis) were collected from wild-type and miR-202-/-fish. Embryos were collected at different stages (1-cell, 8-cell, stage 11/12, stage 17/18, stage 25/26, stage 29/30 and stage 39), according to the developmental table described by Iwamatsu *et al*. (59). For tissues/organs dissection, adult medaka fishes were euthanized by immersion in a lethal dose of tricaine à 30-50mg/L. All tissues/organs and embryos were immediately frozen in liquid nitrogen and subsequently stored at −80°C until RNA extraction. For histological analyses, ovaries were collected from wild-type females, fixed overnight in 4% paraformaldehyde (PFA) at 4°C and dehydrated in 100% methanol and stored at −20°C, before histological analyses.

### Establishment of the *miR-202* mutant medaka line

For the CRISPR/Cas9 knock-out analysis, the target genomic sequence was identified with the help of the ZiFiT online tool (http://zifit.partners.org/ZiFiT/) and using the medaka genome reference available on the Ensembl genome database (Ensembl gene: ENSORLG00000021212). A short sequence in the mature miR-202-3p was selected as followed: GG-(N)18-NN. Two inverse-complementary primers (Forward 5’-TAGGCATAAGGCATGGGAAAAT-3’ and (Reverse 5’-AAACATTTTCCCATGCCTTATG-3’) were annealed and cloned into the pDR274 vector (Addgene plasmid 42250) in the BsaI cloning site. The modified pDR274 vector was digested with DraI and the miR-202 specific guide RNA (mir202-sgRNA) was transcribed using the T7 RNA polymerase (P207, Promega). For the Cas9-RNA *in vitro* synthesis, the pCS2-nCas9n vector (Addgene plasmid 47929) was linearized with NotI and capped RNA encoding the Cas9 was transcribed with the mMessage mMachine SP6 Kit (AM1340, Life Technologies) following manufacturer’s instructions. Cas9 and sgRNA were purified using phenol/chloroform and precipitated by Ammonium acetate. *Cas9*-RNA (100 ng/μL) and *mir202*-sgRNA (10 ng/μL) were co-injected into one-cell stage embryos. Injected embryos were raised to sexual maturity and 10 fishes were genotyped to identified founder fishes (F0). Fishes harboring the same INDEL mutation (−7+3) were selected and outcrossed with wild-type fishes to obtain F1 heterozygous. Such outcrosses were performed at each generation in order to maintain the line. Homozygous fishes were produced for histological and molecular phenotyping analyses. A second line harboring another INDEL mutation (−8) was used in order to confirm the reproductive phenotype, but this line was not used for further histological and molecular analyses.

### Genotyping

Genomic DNA was extracted from a small piece of the caudal fin sampled from anesthetized adult fishes. Samples were lysed in 75mL of lysis buffer containing 1,25 M NaOH and 10 mM EDTA (pH 12) incubated at 90C for 1h and were neutralized with 75mL of neutralization solution containing 2 M Tris-HCl (pH 5). To identify F0 founder fishes, genomic DNA around the expected mutation site was sequenced. For systematic genotyping of individuals of the established line, wild-type and mutant (INDEL −7+3) alleles were specifically detected by HIDI-PCR using specific reverse primers for each allele (S2 Table). The HIDI polymerase (Genaxxon bioscience, M3025.0250) was used with the following PCR conditions: 95°C for 2min; and 40 cycles of 95°C for 20 sec, 57°C for 15 sec and 72°C for 30 sec; and then 72°C for 7 min.

### Total RNA extraction

Frozen tissues were lysed with Precellys Evolution Homogenizer (Ozyme, bertin technologies) in TRI Reagent (TR118, Euromedex) and total RNA was extracted using the “nucleospin RNA” kit (740955, Macherey Nagel).

### Quantitative PCR

For expression analysis of mature miRNA forms, 20 ng of total RNA were reverse-transcribed (RT) using the TaqMan advanced miRNA cDNA Synthesis Kit (A28007, Applied Biosystems). Twenty fmol/µl of an external calibrator cel-miR-39-3p (478293_miR, Life technologies) was added in the first step of the RT-Taqman PCR (polyA step) for 20ng of RNA. The cDNA was diluted (1:5) and universal primers (20x miR-Amp Primer Mix, 100029187, Applied Biosystems) were added in the last step of the RT reaction. The TaqMan QPCR was performed using 5 µl of diluted cDNA, 1 µl TaqMan Advanced miRNA Assay solution (CCU001S, Special Product Custom designed advanced miRNA assay, Life technologies) and 10 µl Fast Advanced Master Mix (4444557, Applied Biosystems) in a total volume of 20 µl. Specific modified probes complementary to *miR-202*-5p and −3p were designed as followed: mir-202-3p 5’_FAM_-AGAGGCATAAGGCATGGGAAAA-3’_Quencher_ and mir-202-5p 5’_FAM_-TTCCTATGCATATACTTCTTTG-3’_Quencher_ (Life technologies). QPCR was performed using the Step One Plus system (Applied Biosystems, USA) with the following conditions: 95°C for 20 sec; and 40 cycles of 95°C for 1 sec and 60°C for 20 sec. The relative expression of miRNA within a sample set was calculated from standard curve using Applied Biosystem StepOne V.2.0 software. All QPCR were performed in duplicates. MiR-26a (hsa-miR-26a-5p, A25576, Thermofisher Scientific) or cel-miR-39-3p was used for normalization.

For mRNA and pri-miR-202 expression analysis, 2µg of total RNA was reverse-transcribed using the Maxima First Strand cDNA Synthesis Kit (K1671, ThermoFisher Scientific). The cDNA was diluted (1:20). The SyberGreen QPCR was performed using 4 µl of diluted cDNA, 5 µl of GoTaq QPCR Master Mix 2x (A600A, Promega) and 100 nM of each primer (S3 Table), in a total volume of 10 µl. The QPCR was performed using the Step One Plus system (Applied Biosystems, Foster City, USA) with the following conditions: 95°C for 2 min; and 40 cycles of 95°C for 15 sec and 60°C for 1 min. Standard curves were generated using five serial cDNA dilutions (from 1:2 to 1:32) of a pool of all samples. The relative abundance of target cDNA was calculated from standard curve using Applied Biosystem StepOne V.2.0 software. All QPCR were performed in triplicates and the *rpl7* gene was used for normalization.

### Microarray analysis

Gene expression profiling was conducted using an Agilent 8×60K microarray as previously described (60). Samples were randomly distributed on the microarray for hybridization. The data were processed using the GeneSpring software (Agilent v.14.5) using gMedianSignal values. The gene expression data was scale normalized and log(2) transformed before the statistical analysis. Corresponding data were deposited in Gene Expression Omnibus (GEO) (https://www.ncbi.nlm.nih.gov/geo/) database under the reference GSE111388. The differences between the groups were analyzed with unpaired t-test after application of minimum two-fold change filter with the significance level of 5 % (p < 0.05) after Benjamini-Hochberg correction.

### Fluorescent *in situ* hybridization and immunostaining

For fluorescent *in situ* hybridization (FISH), fixed ovaries were embedded in paraffin and sections (9 μm thickness) were performed with a microtome (HM355, microm). An anti-sense Locked Nucleic Acid (LNA) oligonucleotide was designed and produced by Exiqon A/S to detect the mature miR-202-3p form. Since medaka and human miR-202-5p sequences are identical, we used the hsa-miR-202-5p miRCURY LNA^TM^ miRNA detection probe to detect the mature medaka miR-202-5p form. A LNA^TM^ Scramble-miR probe (5’-GTGTAACACGTCTATACGCCCA-3’) was used as a negative control. All LNA^TM^ probes were double-DIG labeled at both 5’ and 3’ ends. FISH was performed using the microRNA ISH Buffer Set (FFPE) Hybridization Buffer (ref. 90000, Exiqon), following the manufacturer’s instructions with some modifications. Permeabilisation was performed for 7 min at room temperature using of Proteinase-K (10 mg/ml, P2308 Sigma). LNA probes were used at 20 nM at 53°C (30°C below the RNA Tm °C) for 2 hours. Samples were then incubated overnight at 4°C with a rabbit anti-DIG HRP-conjugate antibody (1:500, Roche). For the GFP detection, a chicken anti-GFP (1/500, ref. ab13970, Abcam) was added at this step. The anti-GFP was first detected with a goat anti-chicken AlexaFluor488-conjugate antibody (1/500, ref. A11039, Life Technologies) for 1 hour at room temperature. Then, the anti-DIG-HRP antibody was detected with the TSA-Cy5 substrate (1:50, TSA™ PLUS Cy5 kit, NEL 745001KT, Perkin Elmer) for 15 min at room temperature. All pictures were taken under SP8 confocal.

### Nuclear staining and image analysis

Fixed ovaries were embedded in paraffin and median sections (7 μm thickness) were performed with a microtome (HM355, microm). Nuclei were stained with DAPI (0,1 µg/ml) at room temperature for 15 minutes in the dark. Median sections were washed 1h in PBS at room temperature. Images of whole sections were acquired with a nanozoomer (HAMAMATSU). For quantitative image analyses, individual oocyte’s area was measured using an image analysis software (Visilog 7.2 for Windows). Based on DAPI staining intensity, the nucleus were segmented then the inner part surround by the nucleus and corresponding to oocytes were individually measured. Pictures were taken under a Nikon AZ100 microscope and DS-Ri1 digital camera.

## ACKNOWLEDGMENTS

We thank the INRA-LPGP fish facility staff, and especially Cecile Duret, for fish rearing and husbandry. We also thank the University of Rennes 1 H2P2 facility for the use of the slide scanning nanozoomer.

## Founding

This work was funded by the TEFOR project (Agence National de la Recherche, ANR-II-INBS-0014, http://www.agence-nationale-recherche.fr/investissements-d-avenir/projets-finances/). VT received the funding. This work has also been supported by the ERA-Net COFASP (COFA) AquaCrispr project (Agence National de la Recherche, ANR-16-COFA-0004, http://www.agence-nationale-recherche.fr/suivi-bilan/editions-2013-et-anterieures/environnement-et-ressources-biologiques/era-net-cofasp-cooperation-in-fisheries-aquaculture-and-seafood-processing/). JB received the funding. The funders had no role in study design, data collection and analysis, decision to publish, or preparation of the manuscript.

## Author Contributions

SG performed histological and molecular analyses, participated in mutant fish phenotyping, data analysis and manuscript writing. JBu participated in histological analyses. AB participated in generating the mutant line and in tissue collection and RNA extraction. LH participated in reproductive phenotype analysis. ALC performed the microarray analysis. JM performed bioinformatics analyses. VT and JBo conceived the study, participated in data analyses and wrote the manuscript. VT supervised and coordinated the study. All authors read and approved the final manuscript.

## Additional information

**Supplementary information accompanies this paper**

### Competing financial interests

The authors declare that they have no competing interests.

### Availability of data and material

The microarray expression data are available from the gene expression omnibus (GEO) database (accession # GSE111388, http://www.ncbi.nlm.nih.gov/geo/).

## Supporting information

**S1 Fig.** FISH on ovarian section performed with a scramble control probe

**S1 Table.** Differentially expressed genes obtained through a microarray analysis

**S2 Table.** Primers used for HIDI-PCR

**S3 Table.** Primers used for QPCR

## References

1. Iwamatsu T, Ohta T, Oshima E, Sakai N. Oogenesis in the Medaka Oryzias latipes : Stages of Oocyte Development : Developmental Biology. Zoolog Sci. 1988;5:353–73.

2. Nishimura T, Tanaka M. Gonadal Development in Fish. Sex Dev. 2014 Sep 8;8(5):252–61.

3. Lubzens E, Young G, Bobe J, Cerdà J. Oogenesis in teleosts: how eggs are formed. Gen Comp Endocrinol. 2010 Feb 1;165(3):367–89.

4. Nakamura S, Kobayashi K, Nishimura T, Higashijima S, Tanaka M. Identification of Germline Stem Cells in the Ovary of the Teleost Medaka. Science. 2010 Jun 18;328(5985):1561–3.

5. Donadeu FX, Schauer SN, Sontakke SD. Involvement of miRNAs in ovarian follicular and luteal development. J Endocrinol. 2012 Jan 12;215(3):323–34.

6. Li X, Wang H, Sheng Y, Wang Z. MicroRNA-224 delays oocyte maturation through targeting Ptx3 in cumulus cells. Mech Dev. 2017;143:20–5.

7. Yao G, Liang M, Liang N, Yin M, Lü M, Lian J, et al. MicroRNA-224 is involved in the regulation of mouse cumulus expansion by targeting Ptx3. Mol Cell Endocrinol. 2014 Jan 25;382(1):244–53.

8. Bizuayehu TT, Babiak I. MicroRNA in Teleost Fish. Genome Biol Evol. 2014 Jul 22;6(8):1911–37.

9. Juanchich A, Le Cam A, Montfort J, Guiguen Y, Bobe J. Identification of Differentially Expressed miRNAs and Their Potential Targets During Fish Ovarian Development. Biol Reprod [Internet]. 2013 May 1 [cited 2017 Jun 11];88(5). Available from: https://academic.oup.com/biolreprod/article/88/5/128,1-11/2514159/Identification-of-Differentially-Expressed-miRNAs

10. Wong QW-L, Sun M-A, Lau S-W, Parsania C, Zhou S, Zhong S, et al. Identification and characterization of a specific 13-miRNA expression signature during follicle activation in the zebrafish ovary. Biol Reprod. 2018 Jan 1;98(1):42–53.

11. Liu J, Luo M, Sheng Y, Hong Q, Cheng H, Zhou R. Dynamic evolution and biogenesis of small RNAs during sex reversal. Sci Rep [Internet]. 2015 May 6;5. Available from: https://www.ncbi.nlm.nih.gov/pmc/articles/PMC4421800/

12. Bizuayehu TT, Lanes CF, Furmanek T, Karlsen BO, Fernandes JM, Johansen SD, et al. Differential expression patterns of conserved miRNAs and isomiRs during Atlantic halibut development. BMC Genomics. 2012 Jan 10;13:11.

13. Jing J, Wu J, Liu W, Xiong S, Ma W, Zhang J, et al. Sex-Biased miRNAs in Gonad and Their Potential Roles for Testis Development in Yellow Catfish. PLoS ONE [Internet]. 2014 Sep 17 [cited 2018 Mar 9];9(9). Available from: https://www.ncbi.nlm.nih.gov/pmc/articles/PMC4168133/

14. Xiao J, Zhong H, Zhou Y, Yu F, Gao Y, Luo Y, et al. Identification and Characterization of MicroRNAs in Ovary and Testis of Nile Tilapia (Oreochromis niloticus) by Using Solexa Sequencing Technology. PLoS ONE [Internet]. 2014 Jan 23 [cited 2018 Mar 9];9(1). Available from: https://www.ncbi.nlm.nih.gov/pmc/articles/PMC3900680/

15. Tao W, Sun L, Shi H, Cheng Y, Jiang D, Fu B, et al. Integrated analysis of miRNA and mRNA expression profiles in tilapia gonads at an early stage of sex differentiation. BMC Genomics [Internet]. 2016 May 4 [cited 2018 Mar 9];17. Available from: https://www.ncbi.nlm.nih.gov/pmc/articles/PMC4855716/

16. Qiu W, Zhu Y, Wu Y, Yuan C, Chen K, Li M. Identification and expression analysis of microRNAs in medaka gonads. Gene. 2018 Mar 10;646:210–6.

17. Lai KP, Li J-W, Tse AC-K, Chan T-F, Wu RS-S. Hypoxia alters steroidogenesis in female marine medaka through miRNAs regulation. Aquat Toxicol Amst Neth. 2016 Mar;172:1–8.

18. Armisen J, Gilchrist MJ, Wilczynska A, Standart N, Miska EA. Abundant and dynamically expressed miRNAs, piRNAs, and other small RNAs in the vertebrate Xenopus tropicalis. Genome Res. 2009 Oct;19(10):1766–75.

19. Bizuayehu TT, Babiak J, Norberg B, Fernandes JMO, Johansen SD, Babiak I. Sex-Biased miRNA Expression in Atlantic Halibut (Hippoglossus hippoglossus) Brain and Gonads. Sex Dev. 2012;6(5):257–66.

20. Landgraf P, Rusu M, Sheridan R, Sewer A, Iovino N, Aravin A, et al. A Mammalian microRNA Expression Atlas Based on Small RNA Library Sequencing. Cell. 2007 Jun 29;129(7):1401–14.

21. Ro S, Song R, Park C, Zheng H, Sanders KM, Yan W. Cloning and expression profiling of small RNAs expressed in the mouse ovary. RNA. 2007 Dec;13(12):2366–80.

22. Bannister SC, Smith CA, Roeszler KN, Doran TJ, Sinclair AH, Tizard MLV. Manipulation of Estrogen Synthesis Alters MIR202* Expression in Embryonic Chicken Gonads. Biol Reprod. 2011 Jan 7;85(1):22–30.

23. Xu L, Guo Q, Chang G, Qiu L, Liu X, Bi Y, et al. Discovery of microRNAs during early spermatogenesis in chicken. PloS One. 2017;12(5):e0177098.

24. Juanchich A, Bardou P, Rué O, Gabillard J-C, Gaspin C, Bobe J, et al. Characterization of an extensive rainbow trout miRNA transcriptome by next generation sequencing. BMC Genomics [Internet]. 2016 Mar 1;17. Available from: http://www.ncbi.nlm.nih.gov/pmc/articles/PMC4774146/

25. Bouchareb A, Le Cam A, Montfort J, Gay S, Nguyen T, Bobe J, et al. Genome-wide identification of novel ovarian-predominant miRNAs: new insights from the medaka (Oryzias latipes). Sci Rep [Internet]. 2017 Jan 10 [cited 2017 Jan 27];7. Available from: http://www.ncbi.nlm.nih.gov/pmc/articles/PMC5223123/

26. Zheng P, Dean J. Oocyte-Specific Genes Affect Folliculogenesis, Fertilization, and Early Development. Semin Reprod Med. 2007 Jul;25(04):243–51.

27. Chen J, Cai T, Zheng C, Lin X, Wang G, Liao S, et al. MicroRNA-202 maintains spermatogonial stem cells by inhibiting cell cycle regulators and RNA binding proteins. Nucleic Acids Res. 2016 Dec 19;gkw1287.

28. Lai EC. Two decades of miRNA biology: lessons and challenges. RNA. 2015 Apr;21(4):675–7.

29. Lin D, Sugawara T, Strauss JF, Clark BJ, Stocco DM, Saenger P, et al. Role of steroidogenic acute regulatory protein in adrenal and gonadal steroidogenesis. Science. 1995 Mar 24;267(5205):1828–31.

30. Stocco DM. StAR protein and the regulation of steroid hormone biosynthesis. Annu Rev Physiol. 2001;63:193–213.

31. Guan G, Sun K, Zhang X, Zhao X, Li M, Yan Y, et al. Developmental tracing of oocyte development in gonadal soma-derived factor deficiency medaka (Oryzias latipes) using a transgenic approach. Mech Dev. 2017;143:53–61.

32. Yan Y-L, Desvignes T, Bremiller R, Wilson C, Dillon D, High S, et al. Gonadal soma controls ovarian follicle proliferation through Gsdf in zebrafish. Dev Dyn Off Publ Am Assoc Anat. 2017 Nov;246(11):925–45.

33. Poon S-K, So W-K, Yu X, Liu L, Ge W. Characterization of inhibin α subunit (inha) in the zebrafish: evidence for a potential feedback loop between the pituitary and ovary. Reproduction. 2009 Jan 10;138(4):709–19.

34. Bertho S, Pasquier J, Pan Q, Le Trionnaire G, Bobe J, Postlethwait JH, et al. Foxl2 and Its Relatives Are Evolutionary Conserved Players in Gonadal Sex Differentiation. Sex Dev Genet Mol Biol Evol Endocrinol Embryol Pathol Sex Determ Differ. 2016;10(3):111–29.

35. Herpin A, Adolfi MC, Nicol B, Hinzmann M, Schmidt C, Klughammer J, et al. Divergent Expression Regulation of Gonad Development Genes in Medaka Shows Incomplete Conservation of the Downstream Regulatory Network of Vertebrate Sex Determination. Mol Biol Evol. 2013 Oct;30(10):2328–46.

36. Nakamoto M, Matsuda M, Wang D-S, Nagahama Y, Shibata N. Molecular cloning and analysis of gonadal expression of Foxl2 in the medaka, Oryzias latipes. Biochem Biophys Res Commun. 2006 May 26;344(1):353–61.

37. Uhlenhaut NH, Jakob S, Anlag K, Eisenberger T, Sekido R, Kress J, et al. Somatic sex reprogramming of adult ovaries to testes by FOXL2 ablation. Cell. 2009 Dec 11;139(6):1130–42.

38. Ottolenghi C, Omari S, Garcia-Ortiz JE, Uda M, Crisponi L, Forabosco A, et al. Foxl2 is required for commitment to ovary differentiation. Hum Mol Genet. 2005 Jul 15;14(14):2053–62.

39. Schmidt D, Ovitt CE, Anlag K, Fehsenfeld S, Gredsted L, Treier A-C, et al. The murine winged-helix transcription factor Foxl2 is required for granulosa cell differentiation and ovary maintenance. Development. 2004 Feb 15;131(4):933–42.

40. Nishimura T, Tanaka M. The Mechanism of Germline Sex Determination in Vertebrates. Biol Reprod [Internet]. 2016 Jul 1 [cited 2017 May 17];95(1). Available from: https://academic.oup.com/biolreprod/article-abstract/95/1/30,1-6/2883593/The-Mechanism-of-Germline-Sex-Determination-in

41. Kurokawa H, Aoki Y, Nakamura S, Ebe Y, Kobayashi D, Tanaka M. Time-lapse analysis reveals different modes of primordial germ cell migration in the medaka Oryzias latipes. Dev Growth Differ. 2006 Apr;48(3):209–21.

42. Hu Q, Zhu Y, Liu Y, Wang N, Chen S. Cloning and characterization of wnt4a gene and evidence for positive selection in half-smooth tongue sole (Cynoglossus semilaevis). Sci Rep [Internet]. 2014 Nov 24 [cited 2018 Mar 7];4. Available from: https://www.ncbi.nlm.nih.gov/pmc/articles/PMC4241513/

43. Charlier C, Montfort J, Chabrol O, Brisard D, Nguyen T, Le Cam A, et al. Oocyte-somatic cells interactions, lessons from evolution. BMC Genomics. 2012 Oct 19;13:560.

44. Faria JAQA, Corrêa NCR, de Andrade C, de Angelis Campos AC, dos Santos Samuel de Almeida R, Rodrigues TS, et al. SET domain-containing Protein 4 (SETD4) is a Newly Identified Cytosolic and Nuclear Lysine Methyltransferase involved in Breast Cancer Cell Proliferation. J Cancer Sci Ther. 2013 Jan 21;5(2):58–65.

45. Dai L, Ye S, Li H-W, Chen D-F, Wang H-L, Jia S-N, et al. SETD4 Regulates Cell Quiescence and Catalyzes the Trimethylation of H4K20 during Diapause Formation in Artemia. Mol Cell Biol [Internet]. 2017 Mar 17 [cited 2018 Mar 7];37(7). Available from: https://www.ncbi.nlm.nih.gov/pmc/articles/PMC5359430/

46. Zhang M, Su Y-Q, Sugiura K, Xia G, Eppig JJ. Granulosa Cell Ligand NPPC and Its Receptor NPR2 Maintain Meiotic Arrest in Mouse Oocytes. Science. 2010 Oct 15;330(6002):366–9.

47. Dickinson RE, Duncan WC. The SLIT/ROBO pathway: a regulator of cell function with implications for the reproductive system. Reprod Camb Engl. 2010 Apr;139(4):697–704.

48. Wong QW-L, Sun M-A, Lau S-W, Parsania C, Zhou S, Zhong S, et al. Identification and characterization of a specific 13-miRNA expression signature during follicle activation in the zebrafish ovary. Biol Reprod. 2017 Dec 6;

49. Zhang J, Liu W, Jin Y, Jia P, Jia K, Yi M. MiR-202-5p is a novel germ plasm-specific microRNA in zebrafish. Sci Rep [Internet]. 2017 Aug 1 [cited 2018 Mar 7];7. Available from: https://www.ncbi.nlm.nih.gov/pmc/articles/PMC5539161/

50. Marlow FL, Mullins MC. Bucky ball functions in Balbiani body assembly and animal-vegetal polarity in the oocyte and follicle cell layer in zebrafish. Dev Biol. 2008 Sep 1;321(1):40–50.

51. Imai T, Saino K, Matsuda M. Mutation of Gonadal soma-derived factor induces medaka XY gonads to undergo ovarian development. Biochem Biophys Res Commun. 2015 Nov 6;467(1):109–14.

52. Guan G, Sun K, Zhang X, Zha X, Li M, Yan Y, et al. Developmental tracing of oocyte development in gonadal soma-derived factor deficiency medaka (Oryzias latipes) using a transgenic approach. Mech Dev [Internet]. [cited 2017 Jan 20]; Available from: //www.sciencedirect.com/science/article/pii/S0925477316301320

53. Li CW, Ge W. Regulation of the activin-inhibin-follistatin system by bone morphogenetic proteins in the zebrafish ovary. Biol Reprod. 2013 Sep;89(3):55.

54. Zhang X, Guan G, Li M, Zhu F, Liu Q, Naruse K, et al. Autosomal gsdf acts as a male sex initiator in the fish medaka. Sci Rep [Internet]. 2016 Jan 27 [cited 2018 Mar 8];6. Available from: https://www.ncbi.nlm.nih.gov/pmc/articles/PMC4728440/

55. Wainwright EN, Jorgensen JS, Kim Y, Truong V, Bagheri-Fam S, Davidson T, et al. SOX9 Regulates MicroRNA miR-202-5p/3p Expression During Mouse Testis Differentiation. Biol Reprod [Internet]. 2013 Aug 1 [cited 2018 Mar 7];89(2). Available from: https://academic.oup.com/biolreprod/article/89/2/34,1-12/2514043

56. Kidder GM, Vanderhyden BC. Bidirectional communication between oocytes and follicle cells: ensuring oocyte developmental competence. Can J Physiol Pharmacol. 2010 Apr;88(4):399–413.

57. Zuccotti M, Merico V, Cecconi S, Redi CA, Garagna S. What does it take to make a developmentally competent mammalian egg? Hum Reprod Update. 2011 Jul 1;17(4):525–40.

58. Robinson DN, Cant K, Cooley L. Morphogenesis of Drosophila ovarian ring canals. Dev Camb Engl. 1994 Jul;120(7):2015–25.

59. Iwamatsu T. Stages of normal development in the medaka Oryzias latipes. Mech Dev. 2004 Jul;121(7–8):605–18.

60. Żarski D, Nguyen T, Le Cam A, Montfort J, Dutto G, Vidal MO, et al. Transcriptomic Profiling of Egg Quality in Sea Bass (Dicentrarchus labrax) Sheds Light on Genes Involved in Ubiquitination and Translation. Mar Biotechnol N Y N. 2017;19(1):102–15.

