## Supplementary figures and images for "MicroRNA-202 *(miR-202)* controls female fecundity by regulating medaka oogenesis"

### Supplementary Materials

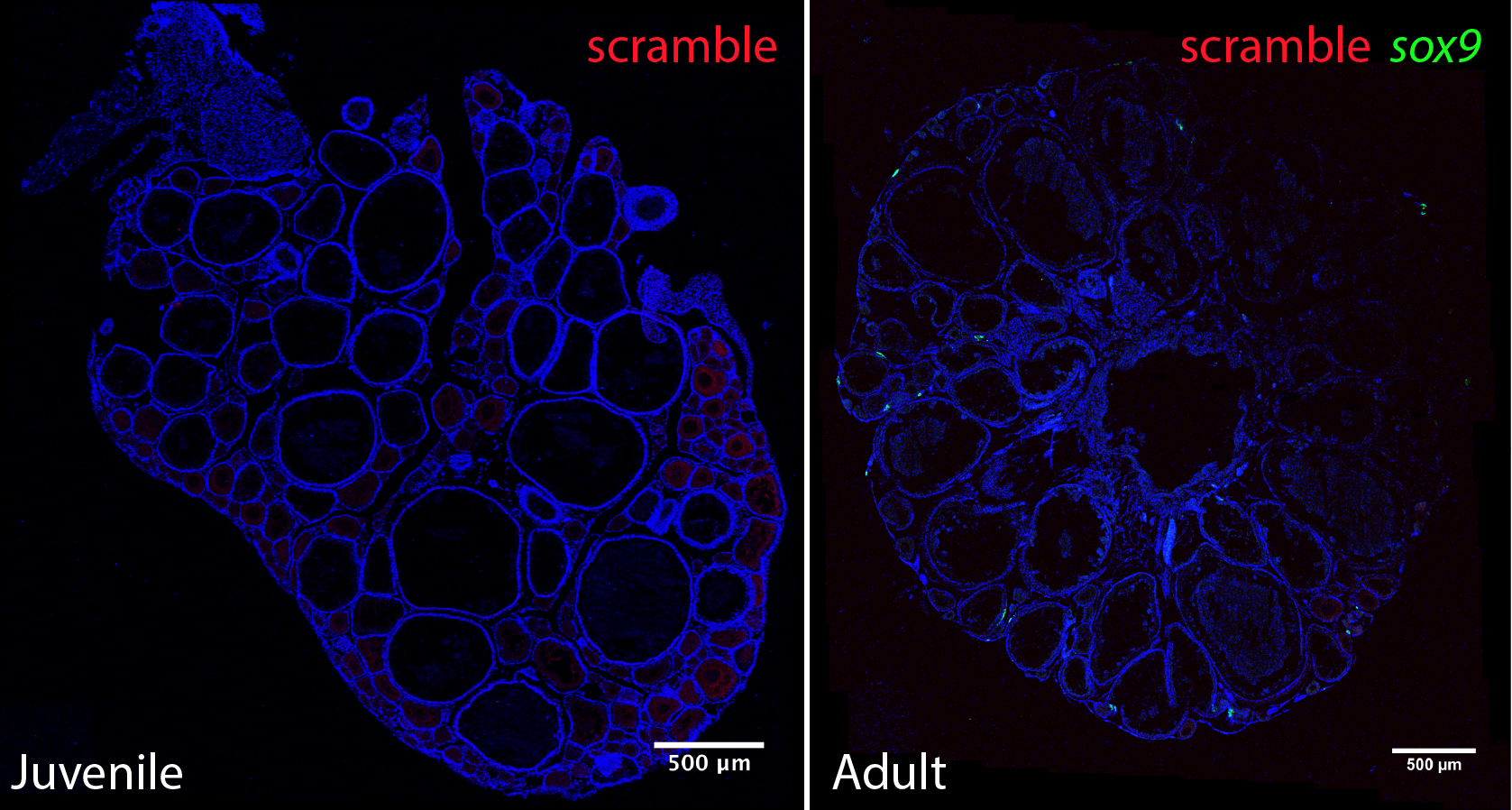
