## Supplementary Materials for "MicroRNA-202 *(miR-202)* controls female fecundity by regulating medaka oogenesis"

S2 Table. Primers used for HIDI-PCR

| Name | Sequence (5’-->3’) |
| --- | --- |
| Genot_HIDI_MiR202_F | CAACCAGTCAATGCACATGAT |
| Genot_HIDI_MiR202WT_R | ACCTCTGCAGCCCCATTTTC |
| Genot_HIDI_MiR202Mut_R | AAAACCTCTGCAGCCCATG |
