## Supplementary Materials for "MicroRNA-202 *(miR-202)* controls female fecundity by regulating medaka oogenesis"

| Gene name | Sequence (5’ --> 3’) |
| --- | --- |
| RPL7_F | GAGATCCGCCTGGCTCGTA |
| RPL7_R | GGGCTGACTCCGTTGATACCT |
| Wnt2b_F | GCACCAGTTCAGACACCATC |
| Wnt2b_R | TGCTTCTCGACTGCTTCTGA |
| Wnt4_F | AGTGTCATGGAGTGTCTGGG |
| Wnt4_R | CCTCAGTGGCACCATCAAAC |
| Clockb_F | CGTCAACAACCAGCAGTGAA |
| Clockb_R | AGTGGAGAATTGAGGCAGCT |
| Setd4_F | CTACCTGGGGCCGTTCTTAA |
| Setd4_R | CGCACACCAGAAAGACACAA |
| Klhl23_F | ATGATGGGAAGCTGAGGTCC |
| Klhl23_R | GGGTGCCAGTAGTATCCTCC |
| Npr1_F | TGAAGATGCCACGGTACTGT |
| Npr1_R | AGCAGACACGTGGATCTTCA |
| Srgap3_F | GCACACCACATTCAGCAGAT |
| Srgap3_R | CCATCATGTTCTCGTCGCTG |
| Cyp19a1a_F | CTCTTCCTGGGTGTTCCTGTTG |
| Cyp19a1a_R | GCTGCTGTCTTGTGCCTCTG |
| Cyp17_F | AGTGACACCAGCCTCGGAGA |
| Cyp17_R | GGTCCACTCCTTCTCATCGTG |
| Gsdf_F | GGGCTGGACACTATTCGAGA |
| Gsdf_R | CATGACACAGAGGAGCTGGA |
| Sox9b_F | AGCGACTCCAAGAAGGACGA |
| Sox9b_R | GTCCAGTCGTAGCCCTTCAG |
| Foxl2_F | GTCACAAACCACAACCTGCT |
| Foxl2_R | TTTGGAGCCGTTTGTCATCC |
| Foxl3_F | CAAAGCCCACCTGAGTCATG |
| Foxl3_R | AGAGCCACGTACGAATAGGG |
| Vasa_F | CCCAAAGTGACCTACATC |
| Vasa_R | AAGTTGATGCCCATCTTG |
| Inh_F | CGTTTCCCTTCCAGCCTTC |
| Inh_R | AAGAGCGTTGCGGATGAG |
| Sycp3_F | ACTTTAGTGGCGGGAAGACG |
| Sycp3_R | GCACATTCATCCGCTCCTTC |
| ControlGeneUp_F | ACTCTGGGTCTCATCTGCAC |
| ControlGeneUp_R | GAGTCACAGCAGGTTCAGGA |

**S3 Table. Primers used for QPCR**
